# Mammalian Y RNAs are modified at discrete guanosine residues with N-glycans

**DOI:** 10.1101/787614

**Authors:** Ryan A. Flynn, Benjamin A. H. Smith, Alex G. Johnson, Kayvon Pedram, Benson M. George, Stacy A. Malaker, Karim Majzoub, Jan E. Carette, Carolyn R. Bertozzi

## Abstract

Glycans modify lipids and proteins to mediate inter- and intramolecular interactions across all domains of life. RNA, another multifaceted biopolymer, is not thought to be a major target of glycosylation. Here, we challenge this view with evidence that mammalian cells use RNA as a third scaffold for glycosylation in the secretory pathway. Using a battery of chemical and biochemical approaches, we find that a select group of small noncoding RNAs including Y RNAs are modified with complex, sialylated N-glycans (glycoRNAs). These glycoRNA are present in multiple cell types and mammalian species, both in cultured cells and *in vivo*. Finally, we find that RNA glycosylation depends on the canonical N-glycan biosynthetic machinery within the ER/Golgi luminal spaces. Collectively, these findings suggest the existence of a ubiquitous interface of RNA biology and glycobiology suggesting an expanded role for glycosylation beyond canonical lipid and protein scaffolds.

## MAIN

Glycans have been shown to regulate a wide array of critical biological processes, ranging from cell-cell contacts to host-pathogen interactions, and even the organization of multicellular organisms(*1*). In a traditionally adjacent field of study, RNA represents another biopolymer that is central to all known life. While the building blocks of RNA are canonically limited to four bases, post-transcriptional modifications (PTMs) can dramatically elaborate the chemical diversity of RNA, with >100 identified PTMs(*2–4*). The cellular role for RNA is more complex than that of a simple messenger. For instance, RNAs function as scaffolds, molecular decoys, enzymes, and network regulators across the nucleus and cytosol(*5–7*). With the exception of a few monosaccharide-based tRNA modifications(*8, 9*), there has been no evidence of a direct interface between these two fields of biology.

We previously developed a strategy for unbiased discovery of protein-associated glycans based on metabolic labeling and bioorthogonal chemistry(*10–12*). We metabolically label cells or animals with precursor sugars functionalized with a clickable azide group. Once incorporated into cellular glycans, the azidosugars enable bioorthogonal reaction with a biotin probe for enrichment, identification, and visualization. In the course of performing such experiments using an azide-labeled precursor to sialic acid, peracetylated N-azidoacetylmannosamine (Ac_4_ManNAz), we made the surprising finding that azide reactivity was present on highly purified RNA preparations from labeled cells. Although there is currently no precedent for a connection between sialoglycans and RNA, either direct or indirect, the fact that RNA is so broadly post-transcriptionally modified in cells motivated us to pursue further investigation.

To explore the possible existence of RNA modified with sialylated glycans (hereafter glycoRNA), we labeled HeLa cells, a human immortalized cell line derived from cervical cancer, with 100 μM Ac_4_ManNAz for up to 48 hours and then used a rigorous protocol to chemically and enzymatically extract RNA in high purity (**Fig. 1A**). To visualize azide-modified components, the RNA sample was reacted with dibenzocyclooctyne-biotin (DBCO-biotin) via Cu-free click chemistry(*13*), separated by denaturing gel electrophoresis and analyzed by blotting (**Fig. 1B**). In an Ac_4_ManNAz- and time-dependent manner, we observed biotinylated species in the very high (>10 kilobases) molecular weight (MW) region. It has recently been reported that high doses of azidosugars can produce non-enzymatic protein labeling(*14*), however, *in vitro* incubation of total RNA with up to 20 mM Ac_4_ManNAz did not produce the previously observed biotinylated species on RNAs in the high MW region (**fig. S1A**). Minor background *in vitro* labeling was apparent on the 28S rRNA, which can also be seen more variably in some Ac_4_ManNAz-labeled cellular RNA experiments (e.g. **Fig. 1B**), but no such background labeling was observed in the putative glycoRNA species.

**Figure 1.**
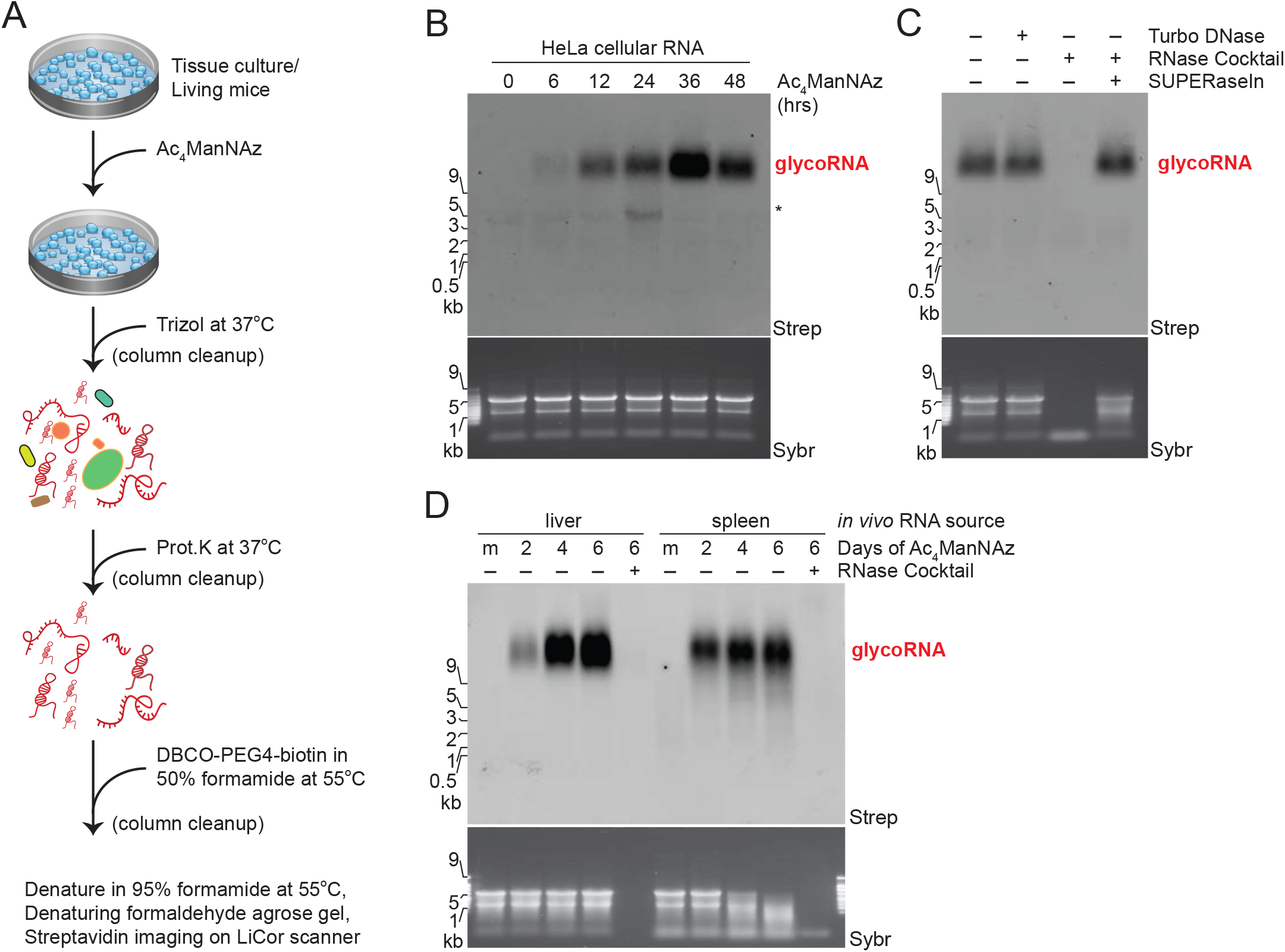
Ac_4_ManNAz, a glycan reporter, incorporates into mammalian cellular RNA. (**A**) Schematic of RNA extraction protocol. Ac_4_ManNAz = peracetylated N-azidoacetylmannosamine. Prot.K = proteinase K. DBCO =dibenzocyclooctyne. (**B**) RNA blotting of RNA from HeLa cells treated with 100 μM Ac_4_ManNAz for the indicated amount of time. After RNA purification, Ac_4_ManNAz was conjugated to DBCO-biotin, visualized with Streptavidin-IR800 (Strep), and imaged on an infrared scanner. Before RNA transfer to the membrane, total RNA was stained and imaged with SybrGold (Sybr) to interrogate quality and loading. All subsequent blots were prepared in this manner, and Ac_4_ManNAz is always used at 100 μM. The regions where glycoRNAs are present (red text) and non-specific labeling (*) is noted. (**C**) RNA Blot of Ac_4_ManNAz-labeled HeLa RNA treated with Turbo DNase or RNase cocktail (A/T1) +/- SUPERaseIn (RNase inhibitor). (**D**) RNA Blot of murine RNA after *in vivo* Ac_4_ManNAz delivery via intraperitoneal injection on indicated days at 300 mg Ac_4_ManNAz/kg/day. RNA from the liver and spleen were analyzed. Mock (m) mice were injected with DMSO only. RNase treatment was performed on extracted RNA.

Further, treatment of RNA from Ac_4_ManNAz-labeled HeLa cells with DNase did not affect the glycoRNA signal while treatment with an RNase cocktail (A and T1) efficiently digested the total RNA and as well as the biotinylated glycoRNA (**Fig. 1C**). This effect required RNase enzymatic activity as pre-blocking of the RNases with an inhibitor, SUPERaseIn, completely rescued the biotinylated glycoRNA (**Fig. 1C**). GlycoRNA was also sensitive to exonucleases, such as RNaseR (3’-5’ exo) and Terminator nuclease (5’-3’ exo), in a manner proportional to the amount of total RNA each enzyme was able to degrade (**fig. S1B**). Thus, cells treated with Ac_4_ManNAz enzymatically incorporate the azide label into cellular RNA which migrates as high MW species.

Using the same metabolic labeling approach, we looked for the presence of glycoRNA in other cell types and in animals. Human embryonic stem cells (H9), a human myelogenous leukemia line (K562), a human lymphoblastoid cell line (GM12878), a mouse T-cell acute lymphoblastic leukemia cell line (MYC T-ALL 4188), and a hamster Chinese hamster ovary cell line (CHO) all showed evidence of the presence of glycoRNA (**figs. S1C and S1D**). In particular, H9 and 4188 cells in particular showed significantly more labeling with Ac_4_ManNAz per mass of total RNA than other cell types (**figs. S1C and S1D**). Next, we assessed if this labeling could occur *in vivo*. To this end, we performed intraperitoneal injections of Ac_4_ManNAz into mice for 2, 4, or 6 days(*15*). In the liver and spleen, which were the organs which yielded enough total RNA for analysis, we observed dose-dependent and RNase-sensitive Ac_4_ManNAz labeling of RNAs in the same MW region as glycoRNAs from cultured cells (**Fig. 1D**). These data suggest that glycoRNA is not an artifact of tissue culture and occurs broadly across multiple cell and tissue types, albeit at various abundances.

Across all cell types and organs tested, glycoRNA was found to migrate very slowly by denaturing agarose gel electrophoresis (**Fig. 1**). We hypothesized that, if glycoRNAs are indeed large RNAs, they would likely be polyadenylated (poly-A). However, we were consistently unable to purify glycoRNA from extracted RNA via poly-A enrichment (**Fig. 2A**). This was not due to cleavage of the glycoRNA during the poly-A enrichment procedure (**fig. S2A**). To address this, we used a fractionation strategy that leverages length-dependent RNA precipitation and binding to silica columns to separate out “large” (>200 nts) from “small” (<200 nts) transcripts. To our surprise, the glycoRNA fractionated exclusively with the small RNA population of the total RNA (**Fig. 2B**). To validate this observation with an independent fractionation strategy, we applied Ac_4_ManNAz-labeled RNA to a sucrose gradient and analyzed the distribution of total RNA via SybrGold staining and glycoRNA. The sucrose gradient efficiently separated the major visible RNAs such as small RNAs/tRNA, 18S rRNA, and 28S rRNA (**Fig. 2C and fig. S2B**). The glycoRNA signal was specifically present in the small RNA fractions, but still demonstrated extremely slow migration (high apparent MW) in the agarose gel (**Fig. 2C**).

**Figure 2.**
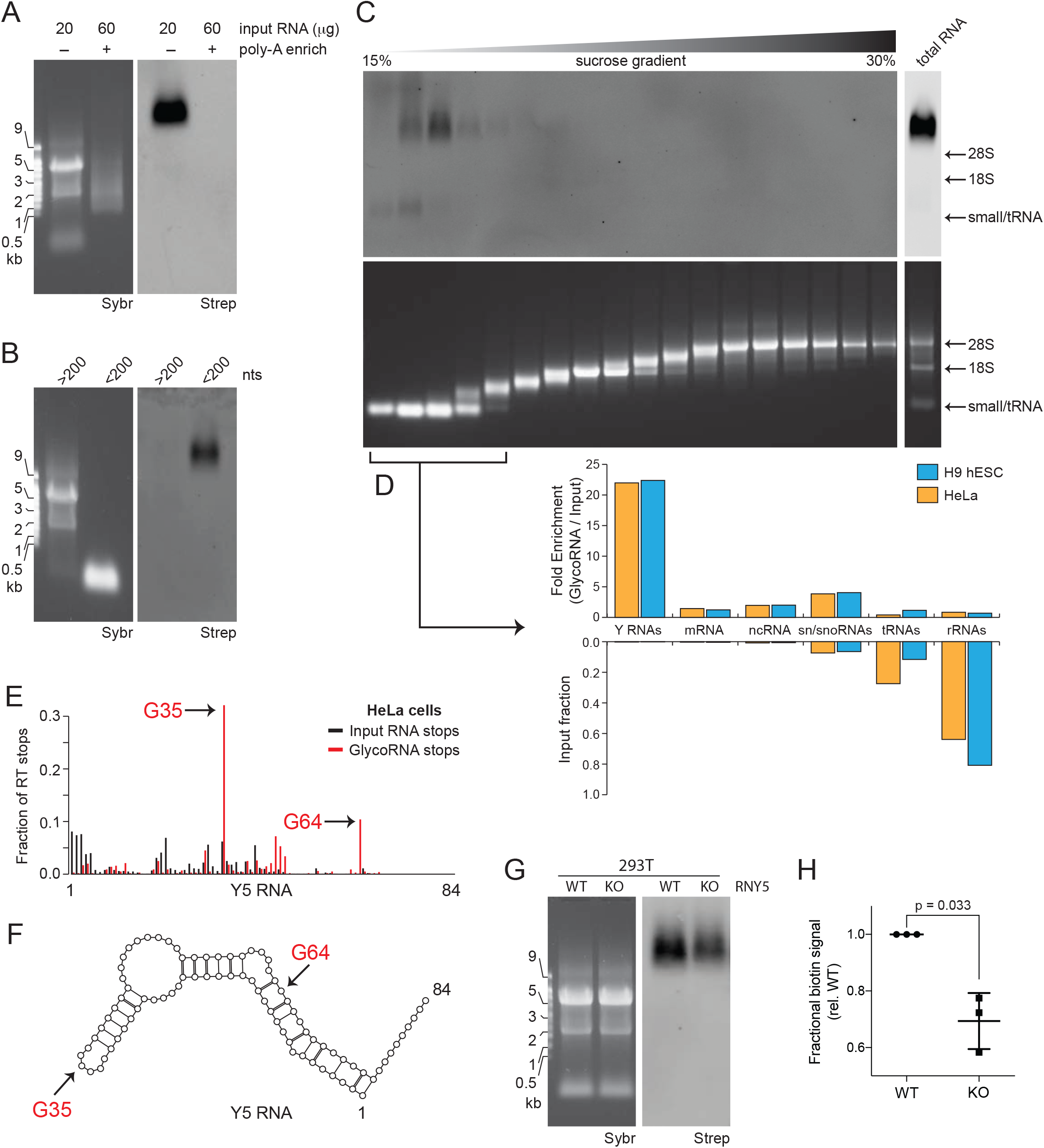
Small, non-polyadenylated, and conserved transcripts comprise the pool of cellular glycoRNA. (**A**) Blotting of total or poly-adenylated (poly-A) enriched RNA from HeLa cells treated with Ac_4_ManNAz. (**B**) Blotting of total RNA from HeLa cells treated with Ac_4_ManNAz after differential precipitation fractionation using silica-based columns. (**C**) Blotting of total RNA from H9 cells treated with Ac_4_ManNAz after sucrose density gradient (15-30% sucrose) fractionation. An input profile is displayed to the right of the gradient. (**D**) Small RNAs were isolated after sucrose density gradients from HeLa (orange) and H9 (blue) cells treated with Ac_4_ManNAz, conjugated to biotin, enriched with streptavidin beads, and sequenced. Input levels of each RNA type are displayed in the bottom bar plot, and fold enrichment (Ac_4_ManNAz reads/input reads) are displayed in the top bar plot. (**E**) Histogram of reverse transcriptase (RT) stops mapped to the Y5 RNA transcript; x-axis is the fraction of RT stops and y-axis is the RNA length. RT stops from the input (black) and Ac_4_ManNAz-enriched (red) libraries from HeLa cells are shown. Two prominent RT stops in the Ac_4_ManNAz-were enriched: Guanosine 35 (G35) and G64. (**F**) Secondary structure of Y5 RNA with the positions of G35 and G64 annotated. Base-pairs shown with lines and nucleotides with open circles. (**G**) Representative blot of total RNA from wild-type (WT) or Y5 knockout (KO) 293T cells treated with Ac_4_ManNAz. (**H**) Quantification of the blot in (G) from biological triplicates. P value calculated by a paired, two-tailed t test.

To identify the glycosylated transcripts, we leveraged the sucrose gradients to isolate only the small RNA fractions from Ac_4_ManNAz-labeled H9 and HeLa cells. RNA sequencing libraries were generated from small RNAs (input) as well as glycoRNAs that were enriched after streptavidin pulldown. As expected, the distribution of RNA transcripts was biased for small non-polyadenylated RNAs (**Fig. 2D, fig. S3A, and table S1**). Quality controls revealed that biological replicates had high concordance, with glycoRNA-enriched and input samples clustering away from each other (**fig. S3B and S3C**). For each small RNA family, an enrichment was calculated to assess the relative distribution of reads in the glycoRNA-enriched vs input samples (**Fig. 2D**). Ribosomal RNA and tRNAs were depleted or poorly enriched, except for the 5.8S rRNA, in the glycoRNA pool (**Fig. 2D**). Messenger RNAs and other noncoding RNA transcripts were poorly enriched (ranging from 1.2–2x over input). However, small nuclear and small nucleolar RNAs (sn/snoRNAs) were the second most enriched class at 3.8-4x over input (**Fig. 2D**). Finally, the Y RNA family stood out as the single best represented set of small RNAs at approximately 22-fold enriched over input (**Fig. 2D**). Notably, despite the differences in cell type and culture conditions, HeLa and H9 glycoRNA enrichment results were highly similar (**Fig. 2D**).

Most snRNAs and snoRNAs were enriched to some degree, with U3, U6, and U8 ranking among the most highly enriched in both HeLa and H9 cells (**fig. S3D**). Among the Y RNAs, Y5 was the most well represented (**fig. S3D**). Finally, considering all the individual snoRNAs, we found a positive correlation between the enrichment of these transcripts across cell types (**fig. S3E**). Exceptions to this were the SNORD116 snoRNAs that are not expressed in HeLa but are expressed and enriched in H9 cells (**table S2**).

Next, we identified putative sites of glycoRNA modification by mapping of reverse transcriptase (RT) stops. RTs are characterized to frequently stop one nucleobase 3’ of PTMs in RNA transcripts((*16, 17*), **Methods**). We chose to focus on Y5 and U3 as they were both highly enriched from HeLa and H9 cells and comprised approximately 3% and 10%, respectively, of all reads in the glycoRNA-enriched libraries (**table S1**). We found that input RNA RT stops were broadly distributed and partially biased for the 5’ ends, which would represent cDNA synthesis templated by an intact and unmodified RNA (**Fig. 2E, fig. S3F, S3G, and table S3**). In contrast, glycoRNA RT stops were less biased for the full-length RNA. This pattern was particularly pronounced for the Y5 glycoRNA, which yielded two strong RT stops at single stranded guanosines 35 (G35) and 64 (G64) (**Fig. 2E, 2F**). Strikingly, the positions and relative intensities were conserved among glycoRNAs from HeLa and H9 cells (**Fig. 2E and fig. S3F**). The RT stop pattern for U3 glycoRNA was more complex, but many of the strongest RT stops occurred at guanosine residues (**fig. S3G**). To understand if guanosine modification is a general feature of glycoRNAs, we calculated the fractional distribution of nucleobases at RT stops from input and enriched libraries along snRNA and Y RNA transcripts. In both cases, guanosine was the only enriched nucleobase (**fig. S3H**).

The RNAs we found to be modified have many well-established and critical cellular roles. The Y RNA family stood out as the most strongly enriched family in our dataset and is highly conserved in vertebrates(*18*). Y RNAs are thought to contribute to cytosolic ribonucleoprotein (RNP) surveillance, particularly for the 5S rRNA(*18, 19*). Additionally, they are (among other glycoRNAs we identified) known to be antigens associated with autoimmune diseases, such as systemic lupus erythematosus and mixed connective tissue disease(*20*). Given these features, we sought to validate Y5 as a glycoRNA by gene knockout via CRISPR/Cas9. A 293T Y5 knockout cell line was generated using two single-guide RNAs (sgRNAs) that targeted the 5’ and 3’ regions of the Y5 genomic locus (**fig. S4A**). Single cell clones were isolated, and a KO was selected for characterization; PCR amplification of the Y5 locus yielded two amplicons corresponding to two different insertion/deletions (**fig. S4B and S4C**). The KO synthesized no observable Y5 transcript but had no gross growth defects (**fig. S4D and S4E**). Ac_4_ManNAz-labeling of the Y5 KO cells resulted in a significant (~30%) decrease in the amount of biotin signal compared to WT cells, without any apparent MW changes (**Fig. 2G and 2H**). The reduction of glycoRNA signal is consistent with the sequencing data, which identifies Y5 as a major, but not exclusive glycoRNA species.

Next we sought to define the glycan structures on glycoRNAs. The major pathway for Ac_4_ManNAz metabolism in human cells entails conversion to sialic acid, then to CMP-sialic acid, and finally addition to an underlying glycan(*21*). To exclude the possibility that Ac_4_ManNAz is shunted into unexpected metabolic pathways, we used 9-azido sialic acid (9Az-sialic acid), which is directly converted into CMP-sialic acid(*22*) as a metabolic label. Consistent with Ac_4_ManNAz labeling, 9Az-sialic acid produced a similar time-dependent labeling of slowly migrating cellular RNA (**Fig. 3A**). Additionally, treatment of Ac_4_ManNAz-labeled cellular RNA with *Vibrio cholerae* sialidase (VC-Sia) completely abolished the biotin signal without impacting the integrity of the RNA sample, while a heat-inactivated (HI) VC-Sia was unable to reduce the signal (**Fig. 3B**). We assessed the contribution of canonical sialic acid biosynthesis enzymes through the use of P-3F_AX_-Neu5Ac, a cell-permeable metabolic inhibitor of sialoside biosynthesis(*23*). Treatment of HeLa cells with P-3F_AX_-Neu5Ac resulted in a dose-dependent reduction in total glycoRNA signal and a concomitant shift towards higher apparent MW on the blot (**Fig. 3C**). This reduced mobility (appearing higher in the gel) of the glycoRNA likely results from less sialic acid, and thus less negative charge, per glycoRNA molecule.

**Figure 3.**
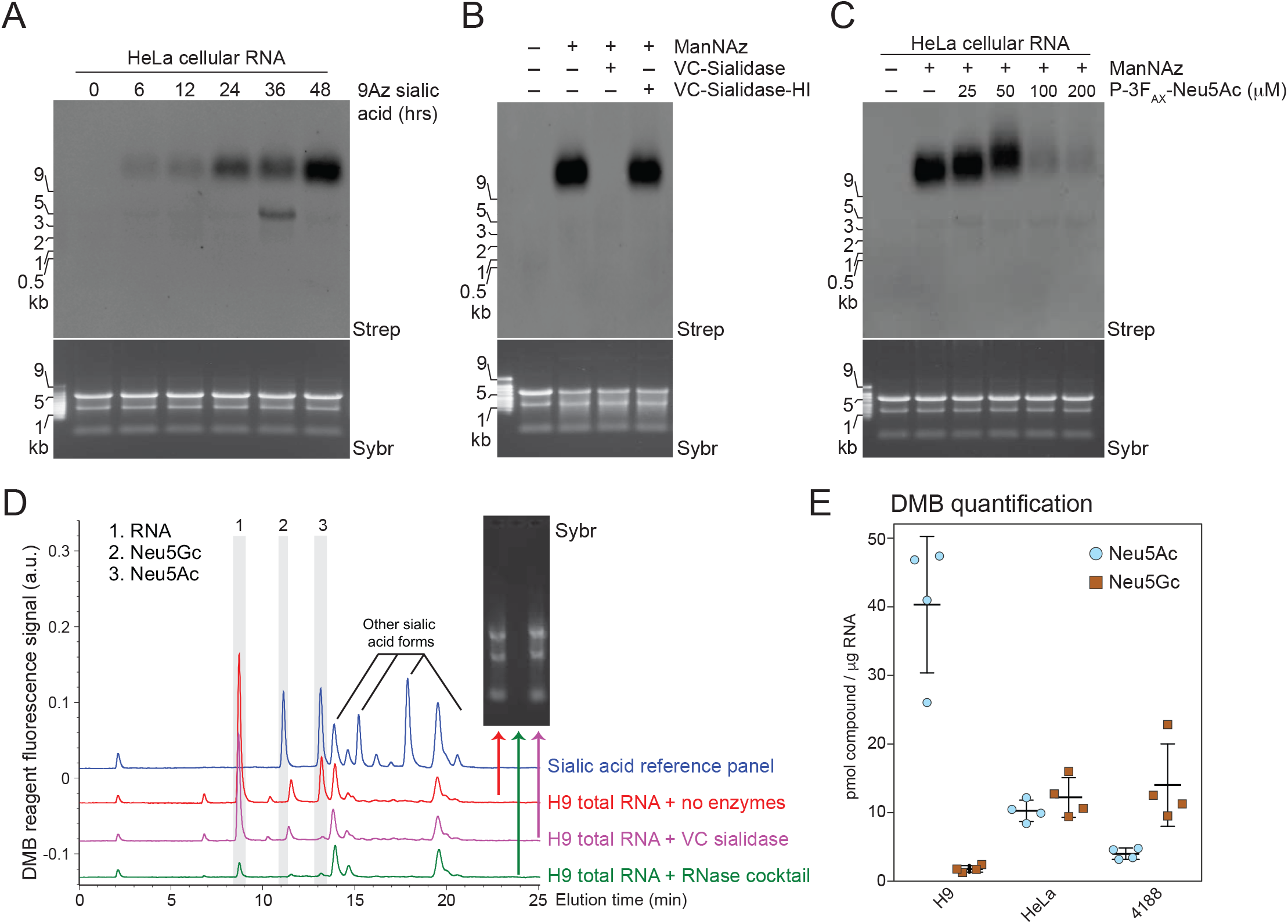
Glycans modifying RNA contain sialic acid. (**A**) Blotting of RNA from HeLa cells treated with 1.75 mM 9-azido sialic acid for indicated times. (**B**) Blotting of Ac_4_ManNAz-labeled HeLa cell RNA treated with *Vibrio cholerae* (VC) Sialidase or Heat-inactivated Sialidase (VC-Sialidase-HI). (**C**) HeLa cells were treated with Ac_4_ManNAz and the indicated concentrations of P-3F_AX_-Neu5Ac. (**D**) Unlabeled total RNA from H9 hESCs (H9) were processed with the fluorogenic 1,2-diamino-4,5-methylenedioxybenzene (DMB) probe and analyzed by HPLC to determine the presence and abundance of specific sialic acids. Inset is a Sybr gel image of the total RNA for each condition: no enzymes, RNase cocktail, or Sialidase treatment. The main sialic acid peaks are #2 and 3. The precise identity of peak 1 is unknown, but it is RNase sensitive. (**E**) Quantification of DMB results from 4188, H9, and HeLa cells from four biological replicates.

To confirm that glycoRNAs are sialylated, we used an independent method not based on metabolic reporters. The fluorogenic 1,2-diamino-4,5-methylenedioxybenzene (DMB) probe is used to derivatize free sialic acids for detection and quantitation by HPLC-fluorescence(*24*). We subjected native, total RNA from HeLa, H9, and 4188 cells to the DMB labeling procedure (**fig. S5A**) and observed the presence of two forms of sialic acid commonly found in animals, Neu5Ac and Neu5Gc (**Fig. 3D and fig. S5B**). These peaks disappeared when the samples were pretreated with VC-Sia or RNase, reinforcing the notion that glycoRNA is modified with sialic acid containing glycans. Notably, we were unable to detect any sialic acid liberated from genomic DNA using the DMB assay (**fig. S5C**).

Quantitatively, we found that H9, HeLa, and 4188 cells have approximately 40, 20, and 20 picomoles (pmol) of total sialic acid per μg of total RNA, respectively (**Fig. 3E**). GlycoRNA from 4188 cells contained more Neu5Gc, whereas H9 cells contained mostly Neu5Ac, and HeLa cells had similar levels of Neu5Ac and Neu5Gc (**Fig. 3E**). Consistently, the DMB results are in line with the observed difference in Ac_4_ManNAz labeling intensity (**fig. S1C and S1D**). Human cells lack a functional *CMAH* gene which is responsible for converting Neu5Ac to Neu5Gc, while this pathway exists in mouse cells(*25*). Correspondingly, we found higher Neu5Gc levels in glycoRNA from mouse 4188 cells as compared to HeLa or H9 cells (**Fig. 3E**). The presence of Neu5Gc in HeLa glycoRNA likely comes from bovine serum in the growth media; H9 cells were grown in serum-free media.

There are two main classes of glycans on proteins, N- and O-glycans, and both can be sialylated. To determine whether glycoRNA structures were related to glycoprotein-associated glycan structures, we used a combination of genetic, pharmacological and enzymatic methods. The *ldlD* mutant CHO cell line lacks the ability to interconvert UDP-glucose(Glc)/galactose(Gal) and UDP-GlcNAc/GalNAc ((*26*) and **fig. S6A**). Thus, in minimal growth media lacking Gal and GalNAc, glycoproteins from *ldlD* CHO have stunted N- and O-glycans because the cells cannot produce UDP-Gal (required for N-glycan elongation) and UDP-GalNAc (required to initiate O-glycosylation). We observed very little glycoRNA labeling in Ac_4_ManNAz-treated *ldlD* CHO cells (**Fig. 4A**) grown in minimal media. However, supplementation of the media with galactose, but not GalNAc, restored glycoRNA labeling, and supplementation with both galactose and GalNAc further boosted labeling intensity (**Fig. 4A**). This result was reproduced using a human K562(*27*) cell line with a CRISPR-Cas9 targeted knockout of UDP-galactose-4-epimerase (GALE), whose activity is lost in the *ldlD* CHO cell line (**fig. S6B**). These results are similar to observations of glycoprotein labeling in these cell types(*28*), suggesting that glycoRNA glycans are structurally related to those found on proteins.

**Figure 4.**
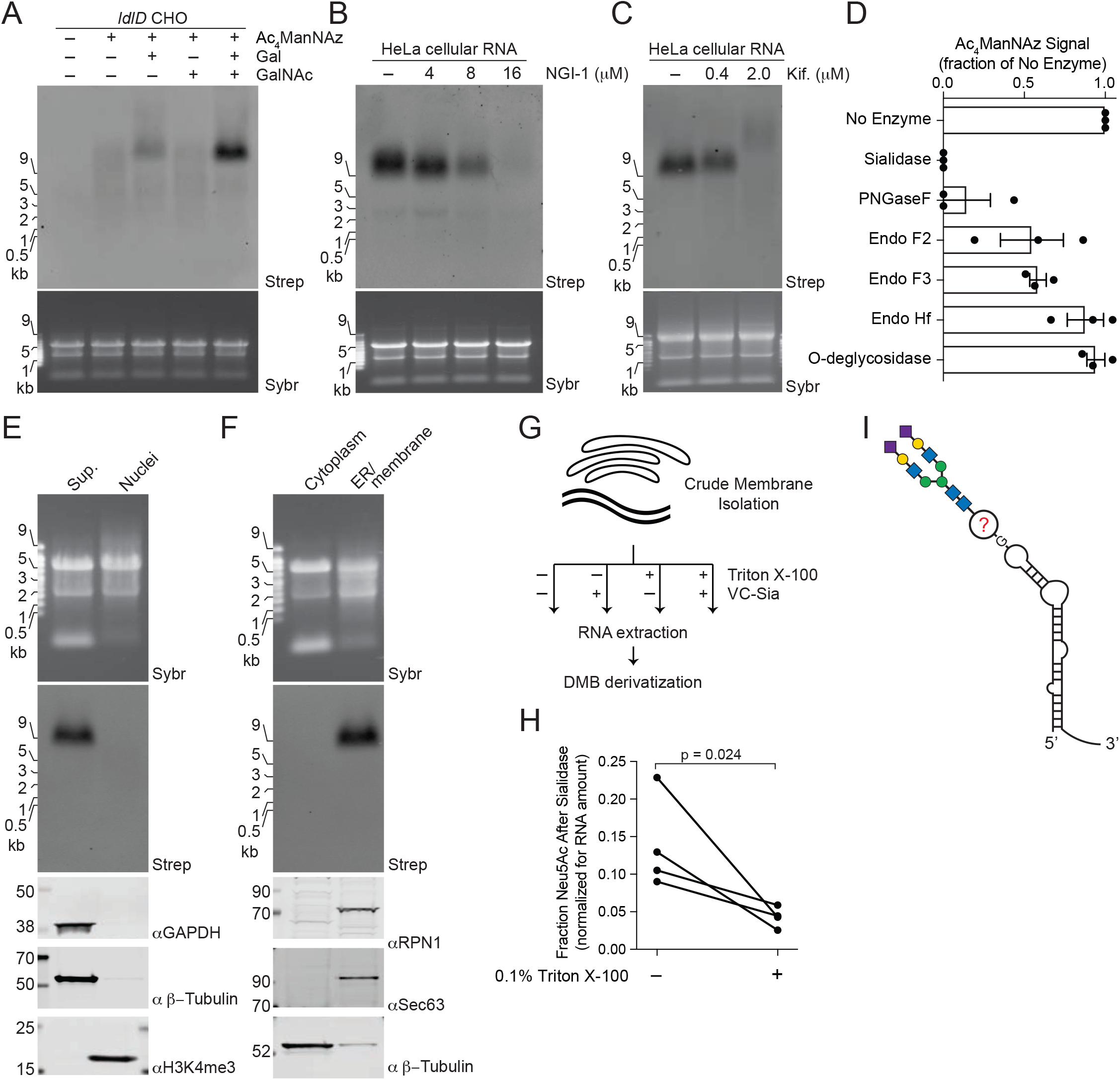
The canonical N-glycan pathway in the ER/Golgi lumen is required for glycoRNA biogenesis. (**A**) Blotting of RNA from *ldlD* CHO cells labeled with Ac_4_ManNAz, Galactose (Gal, 10 μM), N-acetylgalactosamine (GalNAc, 100 μM), or all for 24 hours. (**B**) Blotting of RNA from HeLa cells treated with Ac_4_ManNAz and indicated concentrations of NGI-1, an inhibitor of OST, for 24 hours. (**C**) Blotting as in (B) but with the indicated concentrations of Kifunensine. (**D**) Quantification of Ac_4_ManNAz signal after treatment of Ac_4_ManNAz-labeled HeLa cell RNA with the indicated enzymes *in vitro* each for 1 hour at 37°C in biological triplicate. (**E**) Blotting of RNA and proteins after subcellular fractionation designed to robustly purify nuclei. Non-nuclear proteins GAPDH and β-Tubulin and nuclear Histone 3 lysine 4 trimethylation (H3K4me3) are visualized by western blot. (**F**) Blotting of RNA and proteins after subcellular fractionation designed to robustly separate soluble cytosol from membranous organelles. Membrane proteins RPN1, Sec63, and soluble β-Tubulin are visualized by western blot. (**G**) Schematic of isolated cellular membranes treated with or without Triton X-100 or VC-Sia for 1 hour at 37°C before isolating RNA and assaying sialic acid levels via DMB. (**H**) Quantification of membrane protection assay performed in biological quadruplicates from 293T cells. Significance calculated with an unpaired t-test. (**I**) Model of a glycoRNA: a small RNA is modified at single stranded guanosines with a bi-antennary sialylated N-glycan.

We next tested the effects of glycosylation inhibitors on glycoRNA biosynthesis. Oligosaccharyltransferase (OST) is the major enzyme complex that transfers a 14-sugar precursor lipid-linked oligosaccharide to nascent peptides during their translocation through the Sec/translocon ((*29*) and **fig. S6C**). We tested NGI-1, a specific and potent small molecule inhibitor of OST(*30*), which caused a dose-dependent loss of glycoRNA labeling (**Fig. 4B**), suggesting that OST is involved in biosynthesis of glycoRNA-associated glycans. We also perturbed downstream N-glycan processing steps with kifunensine and swainsonine, inhibitors of the N-glycan trimming enzymes α-mannosidase I and II, respectively ((*31, 32*) and **fig. S6C**), which resulted in a dose-dependent loss of azidosugar labeling (**Fig. 4C and fig. S6D**). This was accompanied by an increased in apparent MW of the glycoRNA at higher doses, akin to the results see with P-3F_AX_-Neu5Ac (**Fig. 3C**). We hypothesize that disruption of high-mannose glycan processing produces hyposialylated glycoRNAs with less net negative charge and, therefore, reduced mobility.

To further define the glycan structures on glycoRNA, we employed a panel of endoglycosidases. Purified RNA from Ac_4_ManNAz-labeled HeLa cells was first exposed to each enzyme and then clicked to biotin for visualization. Treatment of glycoRNA with PNGase F, which cleaves the asparagine side chain amide bond between proteins and N-glycans (*33*), strongly abrogated signal from Ac_4_ManNAz labeling. Endo F2 preferentially cleaves biantennary and high mannose structures, while Endo F3 preferentially cleaves fucosylated bi- and triantennary structures, both within the chitobiose core of the glycan(*33*). Treatment of glycoRNA with either Endo F2 or F3 resulted in a partial loss of Ac_4_ManNAz labeling. However, Endo Hf, which is more specific for high-mannose structures, did not affect Ac_4_ManNAz signal (**Fig. 4D and fig. S6E**). By contrast to these N-glycan digesting enzymes, O-glycosidase (targeting core 1 and core 3 O-glycans(*34*)) or mucinase (StcE (*35*)) treatment had no effect on Ac_4_ManNAz labeling intensity (**Fig. 4D and fig. S6E, F**). As in previous experiments, VC-Sia completely removed the Ac_4_ManNAz signal (**Fig. 4D and fig. S6E**). Together, these data suggest that glycoRNA possesses bi- and tri-antennary N-glycans with at least one terminal sialic acid residue.

Finally, we assessed the subcellular localization of glycoRNA. The biogenesis of sialylated glycans occurs across many subcellular compartments including the cytosol (processing of ManNAc to Neu5Ac), the nucleus (charging of Neu5Ac with CMP), and the secretory pathway (where sialyltransferases append sialic acid to the ends of glycans(*36*)). Interestingly, the localization of Y RNAs has been reported to be mainly cytoplasmic with a minor fraction in the nucleus (*19*). To determine specifically where glycoRNAs are distributed inside cells, we used two biochemical strategies: one which isolates pure nuclei away from membranous organelles mixed with the cytosol(*37*) and a second which separates the soluble cytosolic compartment away from membranous organelles (**Methods**). Nuclear RNA of Ac_4_ManNAz-labeled HeLa cells yielded no detectible azide-labeled species while the ER/membrane fraction quantitatively contained the glycoRNA (**Fig. 4E, F**). This suggests that the glycoRNAs reside within or are closely associated with membrane organelles. To address these possibilities, crude membranes were isolated from Ac_4_ManNAz-labeled 293T cells and subjected to VC-Sia digestion with or without pre-treatment with Triton X-100 to permeabilize membrane organelles (**Fig. 4G**). If glycoRNAs were fully contained within membranes, VC-Sia would only have access to these species after the addition of Triton X-100. We found that the majority of the glycoRNA signal was sensitive to VC-Sia without Triton X-100, while a small but consistent pool was accessible only after permeabilization (**Fig. 4H**). Thus, a proportion of glycoRNAs appears to reside within the luminal space of membranous organelles.

## DISCUSSION

We have found that sialylated N-glycans produced by the canonical ER/Golgi-lumen biosynthetic machinery are attached to specific mammalian small RNAs. These RNAs are a select group of small noncoding RNAs which are consistently modified across several cell types and organisms. Mapping of RT stops suggests that the glycan modifications occur at discrete guanosine residues. Interestingly, the putative sites of glycosylation are predicted to lie within single-stranded loops or bulges (**Fig. 2F**). Overall, these findings point to a common strategy for RNA glycosylation in mammals (**Fig. 4I**).

The glycan-RNA linkage was not sensitive to stringent protocols to separate RNA from lipids and proteins including organic phase separation, proteinase K treatment and silica-based RNA purification. While the precise nature of the glycan-RNA linkage has not yet been determined, we hypothesize that direct glycosylation of native RNA bases is unlikely. The observed sensitivity to PNGase F, which cleaves the glycosidic linkage between Asn and the initiating GlcNAc of N-glycans, implies an amide bond-containing linker that native nucleobases lack. It is possible that a precursor guanosine modification is necessary to establish an asparagine-like functionality capable of modification by OST. Precisely defining the chemical and atomic features of this linkage will be critical for future studies.

Another mystery concerns the mechanism by which small RNA substrates might gain access to luminal compartments in the secretory pathway. There is precedent for transport machineries of intact RNA transcripts across membranes, such as the *C. elegans* transporter SID-1 that imports dsRNA for RNA interference(*38*). Interestingly, mammals have two SID-1 orthologs which are thought to transport RNAs across intracellular membranes(*39, 40*). It is possible that related trafficking systems exist on the ER membrane that would enable N-glycosylation of appropriately functionalized RNA.

Antibodies that cause various autoimmune diseases have been discovered against a striking number of the RNAs, either alone or in complex with proteins, that we identified as glycosylated in this study. A common mechanism by which such RNAs might provoke an immune reaction has been elusive. In light of our findings, the possible access of these RNAs to the secretory pathway and their modification with N-glycan structures may play a role.

The framework in which glycobiology is presently understood excludes RNA as a substrate for N-glycosylation. Our discovery of glycoRNA suggest this is an incomplete view and points to a new axis of RNA glycobiology, including unprecedented enzymology, trafficking, and cell biology.

## Supporting information

Table S1 - Library mapping stats

Table S2 - snoRNA Summary

Table S3 - RT stop histograms

Table S4 - Oligos

## ACKNOWLEDGMENTS

We thank Phillip Sharp, Robert Spitale, Eliezer Calo, Steven Banik, and the Bertozzi Group members for critical discussion. We also thank Hannah Long, Ulla Gerling-Driessen, and Cheen Ang for help with human ES cell culture, Caitlyn Miller for synthesis of 9Az sialic acid, Dean Felsher for the MYC T-ALL 4188 cells, and Melissa Gray for expression and purification of VC-Sialidase. Cell sorting was performed on an instrument in the Shared FACS Facility obtained using NIH S10 Shared Instrument Grant S10RR025518-01.

## Author Contributions

R.A.F. conceived the project. C.R.B. supervised the project. R.A.F., K.P., and C.R.B developed experimental plans. B.A.H.S., S.A.M., and R.A.F. performed and analyzed DMB experiments. A.G.J. and R.A.F. performed and analyzed sucrose gradients. B.M.G. performed mouse related experiments. K.M., R.A.F., and J.E.C. performed CRISRP/Cas9 experiments. R.A.F. and C.R.B. wrote the manuscript. All authors discussed results and revised the manuscript.

## Data and materials availability

All sequencing data has been deposited on the Gene Expression Omnibus (GSE136967).

---

**Supplemental Figure 1.**
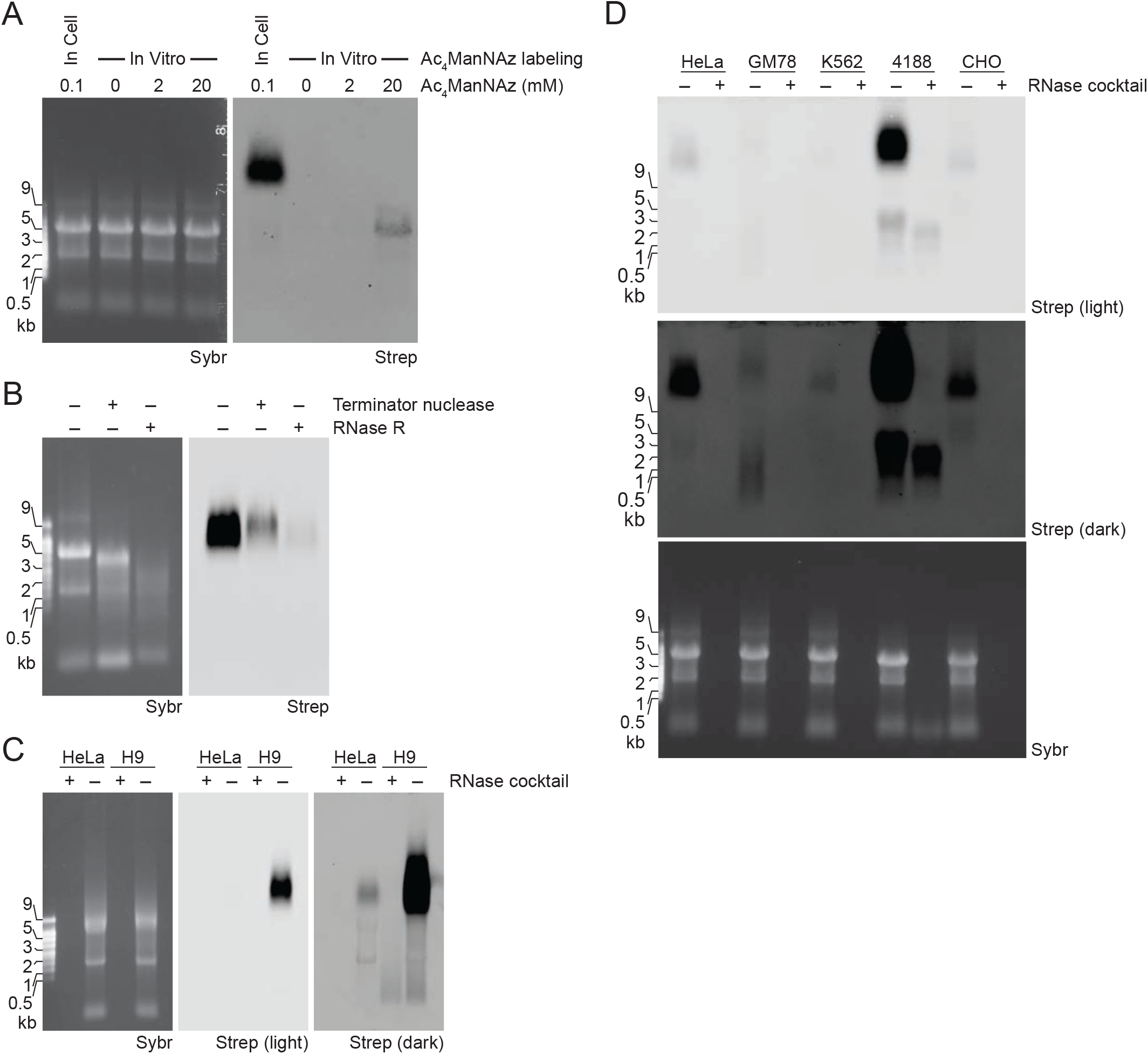
Ac_4_ManNAz incorporation into RNA from multiple mammalian cell types. (**A**) Blotting of *in cellulo* or *in vitro* Ac_4_ManNAz-labeled RNA. Cells were treated with 100 μM Ac_4_ManNAz for 24 hours while native RNA *in vitro* was treated with up to 20 mM Ac_4_ManNAz for 2 hours at 37 °C. (**B**) Blotting of total RNA from Ac_4_ManNAz-labeled HeLa cells treated with no enzymes, Terminator nuclease (a 5’ to 3’ exonuclease), or RNase R (a 3’ to 5’ exonuclease). (**C, D**) Blotting of RNA from various cell types labeled with Ac_4_ManNAz including HeLa and H9 cells (C) and HeLa, GM12878, K562, MYC T-ALL 4188, and CHO cells (D).

**Supplemental Figure 2.**
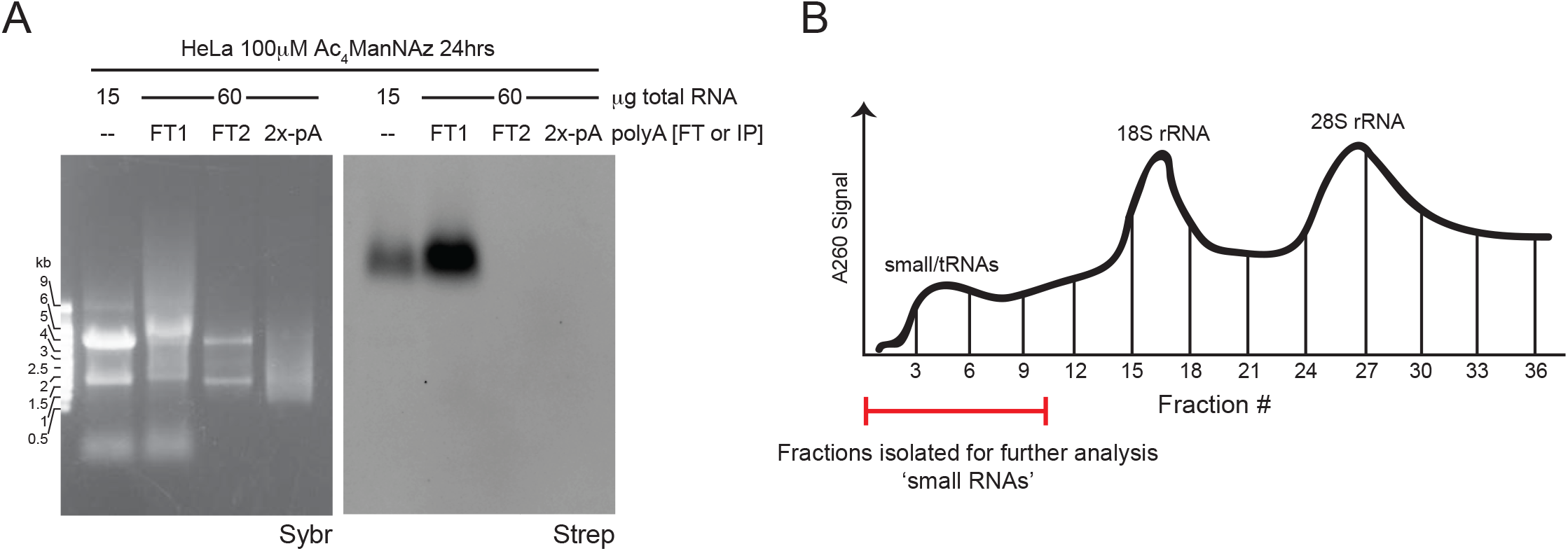
glycoRNAs are non-polyadenylated, small RNAs. (**A**) Blotting of Ac_4_ManNAz-labeled HeLa RNA isolated before and at each step of the poly-A enrichment protocol. Each fraction was collected to assess if specific buffers or temperatures the RNA was exposed to resulted in the loss of Ac_4_ManNAz signal or the degradation of the underlying RNA. Three times the amount of RNA was used for the poly-A enrichment (60 μg) such that even low levels of Ac_4_ManNAz-labeling of poly-A RNA could be detected. FT = flowthrough. 2x-pA = double poly-A enriched RNA. (**B**) Representative A254nm trace of total RNA signal during the fractionation after spinning through a 15-30% sucrose density gradient. Three clear peaks were present representing the small/tRNA RNA, 18S rRNA, and 28S rRNA populations. Fractions containing small RNAs, as indicated on the profile with a red bar, were pooled and purified for sequence analysis.

**Supplemental Figure 3.**
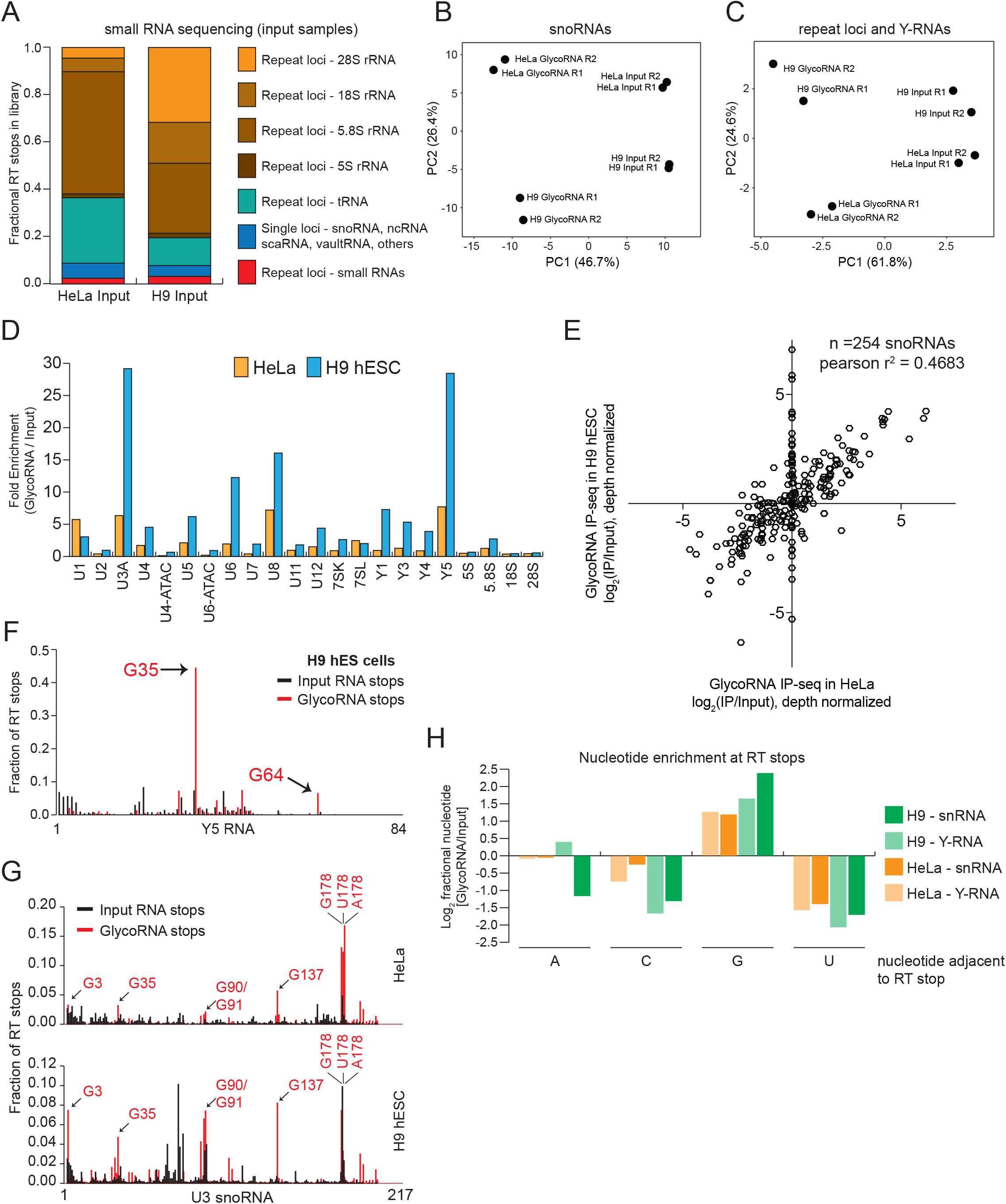
Sequence analysis reveals specific RNAs captured via Ac_4_ManNAz-enrichment. (**A**) Fractional distribution of unique RT stops from HeLa and H9 small RNA sequencing after mapping to the indicated transcripts of the human genome. RNA transcripts that have multiple loci in the genome are labeled as “repetitive” while those with one locus are labeled “single copy”. (**B, C**) Principal component analysis (PCA) of RT stops from input and Ac_4_ManNAz-enriched libraries displaying data from individual biological replicates (total 8 sequencing libraries) after mapping to (B) single copy locus snoRNAs or (C) repeat locus rRNA, snRNA, and single copy Y RNAs. X-axis is the 1^st^ dimension and the y-axis is the second dimension, labeled percentages denote the fractional contribution of each dimension. (**D**) Enrichment analysis of individual repetitive snRNA, snoRNAs, and Y RNAs. Fold enrichment (Ac_4_ManNAz reads/input reads) were calculated for each transcript from HeLa (orange) and H9 (blue) libraries. (**E**) Scatter plot of fold enrichment for 255 snoRNAs in the Ac_4_ManNAz-enrichment. X-axis and Y-axis plot HeLa and H9 vales, respectively. Pearson’s correlation was calculated between the enrichment values for all snoRNAs that had at RT stops in at least one of the datasets (255). (**F**) Histogram of RT stops from H9 cells mapped to the Y5 RNA transcript from the input (black) and Ac_4_ManNAz-enriched (red) libraries. (**G**) Histogram as in (F) mapped to the U3 snoRNA (217 nts long) from HeLa (top) and H9 (bottom) cells. Abundant rT stops in the Ac_4_ManNAz-enriched libraries are highlighted, G = guanosine. (**H**) Nucleotide-level enrichment at RT stops. Ratios of fractional abundance of RT stops from Ac_4_ManNAz-enriched and input libraries were compared in HeLa and H9 libraries across the snRNAs and Y RNA transcripts.

**Supplemental Figure 4.**
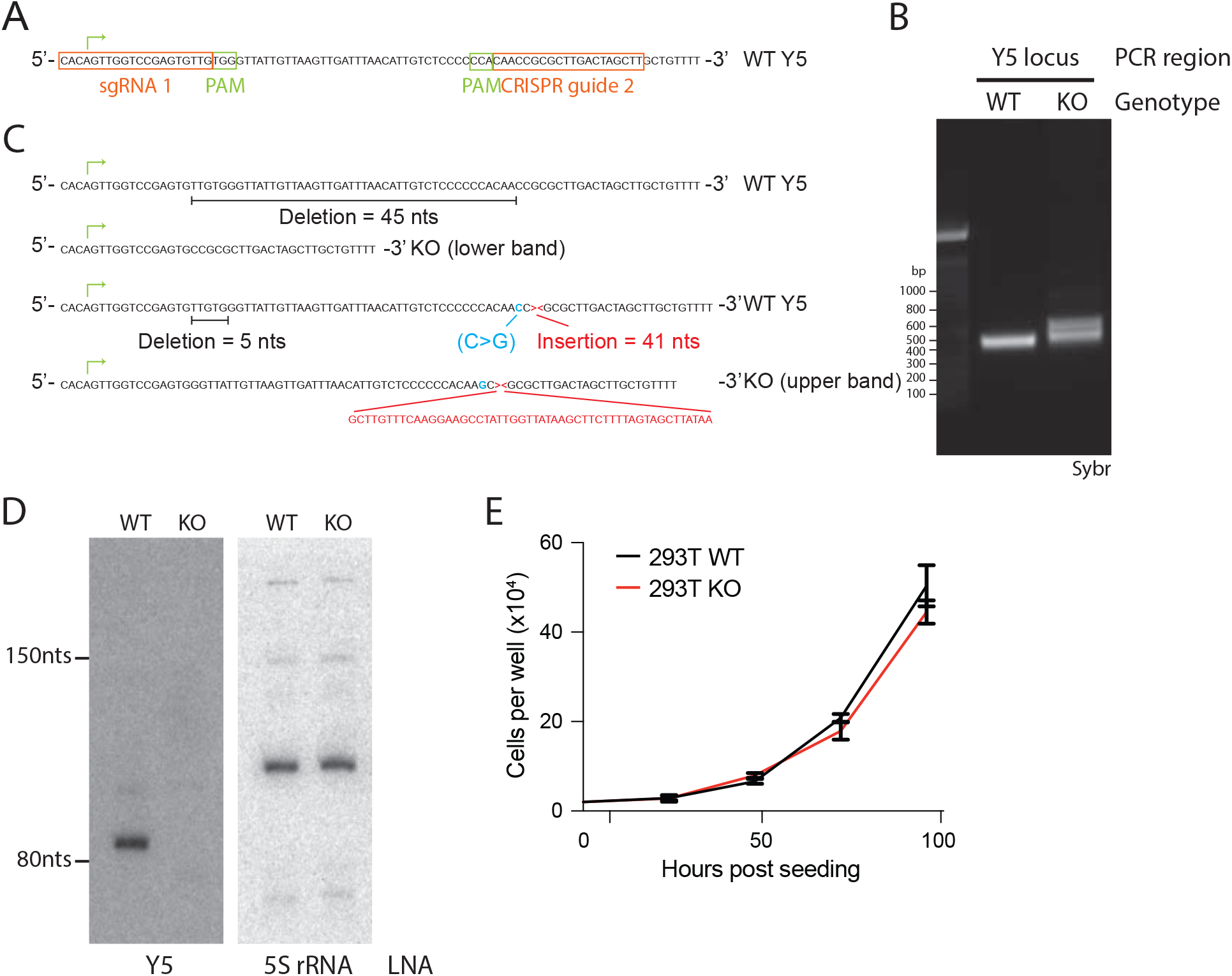
Generation and characterization of Y5 knockout 293T cells. (**A**) Schematic of the wild-type (WT) locus of the human Y5 transcript. The regions targeted by the two designed sgRNAs are highlighted in orange with the protospacer adjacent motif (PAM) regions in green boxes. The first nucleotide of the Y5 RNA is marked with a green arrow. (**B**) PCR analysis of genomic DNA (gDNA) from WT and Y5 KO clone. Primers targeting the Y5 locus (left) showed two new bands in the KO clone. (**C**) Schematic of sanger sequencing results from the gDNA PCR in (B). Two specific sequences were isolated by sequencing. A deletion (black) comes from the bottom band in the gel. A separate insertion (red)/deletion (black)/mutation (blue) structure was observed which comes from the top PCR band. (**D**) Northern blot of 293 WT and KO total RNA. The KO resulted in a complete loss of the Y5 RNA, with the 5S rRNA serving as a loading control. (**E**) Analysis of growth rate between the WT and KO clones across four days of culture. Three independent wells of cells were counted on each day.

**Supplemental Figure 5.**
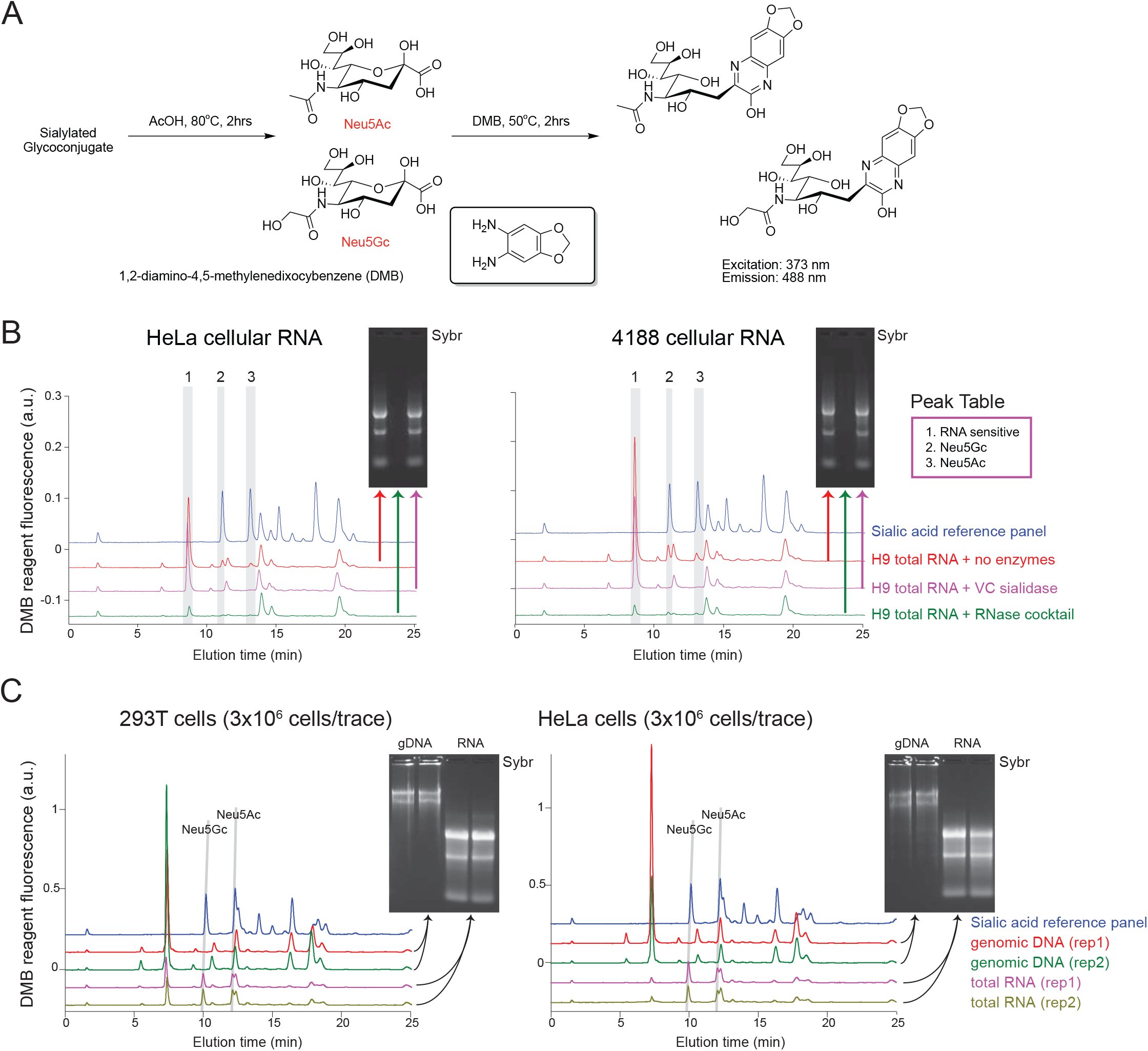
DMB detects and quantifies sialic acids on unlabeled cellular RNA. (**A**) Schematic of experimental steps of the DMB assay with associated structures of the two major types of sialic acid (Neu5Ac and Neu5Gc) as well as the DMB reagent. (**B**) HPLC-fluorescence traces of DMB modified HeLa (left) and 4188 (right) cellular RNA. The main sialic acid peaks are #2 and 3. Peak 1 is unknown but is sensitivity to RNase. Insets are Sybr stained gels of the total RNA for each condition: no enzymes, RNase cocktail, or Sialidase treatment. (**C**) HPLC traces of DMB modified 293T (left) and HeLa (right) cellular RNA or DNA. Neu5Ac and Neu5Gc peaks are highlighted across biological duplicates from each cell type with input material run in a denaturing agarose gel for sizes and loading controls (insets). Only cellular RNA produces Neu5Ac and Neu5Gc specific peaks after DMB derivatization.

**Supplemental Figure 6.**
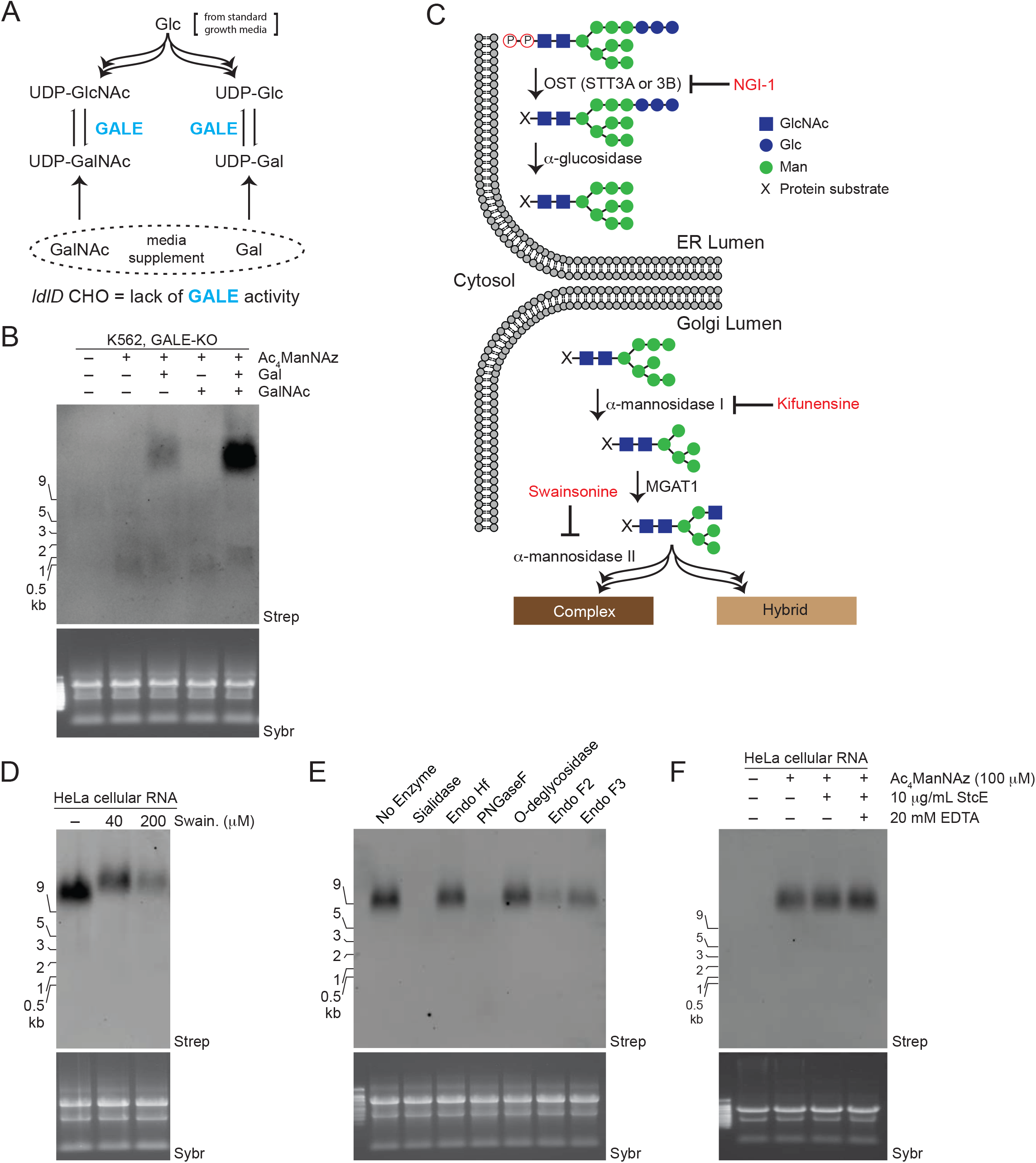
Cellular and *in vitro* assays define the requirement of the N-glycan pathway for glycoRNA. (**A**) Diagram of the main cellular pathway for interconversion of glucose (Glc) to galactose (Gal). Normal cell culture media contains Glu without Gal. The enzyme GALE is responsible for converting uridine 5’-diphospho-N-acetylglucosamine (UDP-GlcNAc) to UDP-GalNAc and UDP-Glc to UDP-Gal. *ldlD* CHO cells contain a loss-of-function mutation in the GALE pathway rendering them deficient in the production of N- and O-linked glycans when grown in normal media. If supplemented with Gal or GalNAc, these cells can selectively elaborate N- and O-linked glycans, respectively. Importantly, biosynthesis of CMP-sialic acid is unaffected by GALE mutation. (**B**) Blotting of RNA from K562-GALE-KO cells labeled with Ac_4_ManNAz, Gal (10 μM), or GalNAc (100 μM) for 24 hours. (**C**) An overview of major steps in the ER- and Golgi-localized steps of N-glycan biosynthesis. Lipid-linked precursor glycans are transferred to ER-lumen-facing peptides (X) by OST. These glycans are sequentially modified by enzymes that first trim and then elaborate the glycan tree. Specific chemical inhibitors such as NGI-1 (inhibiting OST), Kifunesine (inhibiting a-mannosidase I) and Swainsonine (inhibiting α-mannosidase II) are employed to interrogate the importance of each of these enzymes to the accumulation of Ac_4_ManNAz-labeling of RNA. (**D**) Blotting of Ac_4_ManNAz-labeled HeLa cell total RNA after treatment with the indicated concentrations of Swainsonine. (**E**) Blotting of Ac_4_ManNAz-labeled HeLa cell total RNA after *in vitro* treatment with the indicated enzymes; representative of three independent experiments and quantified results are presented in **Fig. 3D**. (**F**) Blotting of Ac_4_ManNAz-labeled HeLa cell total RNA after *in vitro* treatment with the mucinase StcE. Addition of EDTA inhibits the mucinase as a control.

## MATERIALS AND METHODS

### Mammalian cell culture

All cells were grown at 37°C and 5% CO2. HeLa and HEK293T cells were cultured in DMEM media supplemented with 10% fetal bovine serum (FBS) and 1% penicillin/streptomycin (P/S). GM12878, K562, K562^GALE-/-^(*27*), and MYC T-ALL 4188 cells were cultured in RPMI-1640 media with glutamine supplemented with 10% FBS and 1% P/S. CHO and *ldlD*-CHO cells were cultured in 1:1 DMEM:F12 media with 3% FBS and 1% P/S. The H9 human embryonic stem cell line was cultured on Matrigel matrix (Corning) coated plates with mTeSR 1 (StemCell Technologies) media.

### Metabolic chemical reporters and inhibitors

Stocks of azide-labeled sugars N-Acetyl-9-azido-9-deoxy-neuraminic acid (9Az sialic acid, Carbosynth) and N-azidoacetylmannosamine-tetraacylated (Ac_4_ManNAz, Click Chemistry Tools) were made to 500 mM in sterile dimethyl sulfoxide (DMSO). Stocks of unlabeled sugars N-Acetyl-D-galactosamine (GalNAc, Sigma) and D-(+)-Galactose (Gal, Sigma) were made to 500 mM and 50 mM, respectively, in sterile water. In cell experiments ManNAz was used at a final concentration of 100 μM. *In vitro* experiments with ManNAz used 0, 2, or 20 mM ManNAz (up to 200x the in-cell concentrations) for 2 hours at 37°C. The in-cell experiments with 9Az sialic acid used a 1.75 mM final concentration for between 6 and 48 hours. Gal and GalNAc were used as media supplements at 10 μM and 100 μM, respectively, and were added simultaneously with ManNAz for labeling.

Working stocks of glycan-biosynthesis inhibitors were all made in DMSO at the following concentrations and stored at −80°C: 10 mM NGI-1 (Sigma), 10 mM Kifunensine (Kif, Sigma), 10 mM Swainsonine (Swain, Sigma), 50 mM P-3F_AX_-Neu5Ac (Tocris). All compounds were used on cells for 24 hours and added simultaneously with ManNAz for labeling.

### Metabolic reporters in mouse models

All experiments were performed according to guidelines established by the Stanford University Administrative Panel on Laboratory Animal Care. C57Bl/6 mice were crossed and bred in house. ManNAz was prepared by dissolving 100 mg ManNAz in 830 μL 70% DMSO in phosphate buffered saline (PBS), warming to 37°C for 5 minutes, and then sterile filtering using 0.22 μm Ultrafree MC Centrifugal Filter units (Fisher Scientific); this solution was stored at −20°C. Male C57Bl/6 mice (8-12 weeks old) were injected once-daily, intraperitoneally with 100 μL of ManNAz (dosed to 300 mg ManNAz/kg/day), while control mice received the vehicle alone. At 2, 4, and 6 days, mice were euthanized, and their livers and spleens were harvested. The organs were pressed through a nylon cell strainer and resuspended with PBS to create a single cell suspension. RNA was collected as described below.

### RNA extraction and purification strategies

A specific series of steps were taken to ensure that RNA analyzed throughout this study was as pure as possible. First TRIzol reagent (Thermo Fischer Scientific) was used as a first step to lyse and denature cells or tissues. After homogenization in TRIzol by pipetting, samples were incubated at 37°C to further denature non-covalent interactions. Phase separation was initiated by adding 0.2x volumes of 100% chloroform, vortexing to mix, and finally spinning at 12,000x g for 15 minutes at 4°C. The aqueous phase was carefully removed, transferred to a fresh tube and mixed with 2x volumes of 100% ethanol (EtOH). This solution was purified over a Zymo RNA clean and concentrator column (Zymo Research) largely as per the manufacturer’s instructions. Modifications include: (1) each volume of the TRIzol-aqueous phase/EtOH mix was passed over the column twice to fully capture all RNAs and (2) to elute RNAs, two volumes of pure water were used. Next RNA was subjected to protein digestion by adding 1 μg of Proteinase K (PK, Thermo Fischer Scientific) to 25 μg of purified RNA and incubating it at 37°C for 45 minutes. After PK digestion, RNA was purified again with a Zymo RNA clean and concentrator using the protocol as prescribed by the manufacturer. All RNA samples generated in this study were purified at least by these two steps first, with subsequent enzymatic or RNA fractionations occurring in addition to these first two purifications. We found that Zymo-Spin IC and IIICG columns bind up to ~50 and 350 μg of total RNA, respectively; columns in a given experiment were selected based on the amount of RNA needed to be purified.

For differential-precipitation of small vs large RNAs, the Zymo RNA clean and concentrator protocol was used as described. Briefly, RNA in an aqueous solution was mixed with 1x volumes of 50% RNA Binding Buffer in 100% EtOH. This mix was applied to the Zymo silica column; the flow through contained small RNAs while the column retained large RNAs. The flow through was mixed with 1x volumes of 100% EtOH, bound to a new Zymo column and purified as prescribed by the manufacturer.

To enrich for poly-adenylated RNA species, RNA initially purified as above was used as the input for the Poly(A)Purist MAG Kit (Thermo Fisher Scientific). After following the manufacturer’s protocol to elute polyA-tailed transcripts, RNA was subsequently cleaned up via the Zymo RNA clean and concentrator as described above.

### Enzymatic treatment of RNA samples and cells

Various endo- and exonucleases and glycosidases were used to digest RNA, DNA, or glycans. All digestions were performed on 20 μg of total RNA in a 20 μL at 37°C for 60 minutes. To digest RNA the following was used: 1 μL of RNase cocktail (0.5U/μL RNaseA and 20U/μL RNase T1, Thermo Fisher Scientific) with 20 mM Tris-HCl (pH 8.0), 100 mM KCl and 0.1 mM MgCl_2_ or 2 μL of RNaseR (20U/μL, Epicentre) with 20 mM Tris-HCl (pH 8.0), 100 mM KCl and 0.1 mM MgCl_2_ or 2 μL of Terminator nuclease (1U/μL, Epicentre) with 50 mM Tris-HCl (pH 8.0), 2 mM MgCl_2_, 100 mM NaCl. To block the RNase activity of the RNase Cocktail, 1μL of RNase Cocktail was premixed with 8 μL of SUPERaseIn (20U/μL, Thermo Fisher Scientific) for 15 minutes at 25°C before adding to the RNA solution. To digest DNA, 2 μL of TURBO DNase (2U/μL, Thermo Fisher Scientific) with 1x TURBO DNase buffer (composition not provided by manufacture). To digest glycans: 2 μL of α2-3,6,8 Neuraminidase (50U/μL, New England Biolabs, NEB) with GlycoBuffer 1 (NEB), or 2 μL of Endo-Hf (1,000U/μL, NEB) with GlycoBuffer 3 (NEB), or 2 μL of PNGase F (500U/μL, NEB) with GlycoBuffer 2 (NEB), or 2 μL of Endo-F2 (8U/μL, NEB) with GlycoBuffer 3 (NEB), or 2 μL of Endo-F3 (8U/μL, NEB) with GlycoBuffer 4 (NEB), or 2 μL of O-Glycosidase (40,000U/μL, NEB) with GlycoBuffer 2 (NEB), or 1 μL of StcE(*35*) at 0.5 μg/μL with or without 20 mM EDTA. For live cell treatments, VC-Sia was expressed and purified as previously described(*41*) and added to cells at 150 nM final concentration in complete growth media for 1 hour at 37°C.

### Strain-promoted Alkyne-Azide Cycloaddition (SPAAC) conjugation to RNA

Strain-promoted Alkyne-Azide Cycloaddition (SPAAC) was used in all experiments to avoid copper in solution during the conjugate of biotin to the azido sugars (ManNAz and 9Az-Sia). All experiments used dibenzocyclooctyne-PEG4-biotin (DBCO-biotin, Sigma) as the alkyne half of the cycloaddition. To perform the SPAAC, RNA in pure water was mixed with 1x volumes of “dye-free” Gel Loading Buffer II (df-GLBII, 95% Formamide, 18mM EDTA, and 0.025% SDS) and 500 μM DBCO-biotin. Typically, these reactions were 10 μL df-GLBII, 9 μL RNA, 1 μL 10mM stock of the DBCO reagent. Samples were conjugated at 55°C for 10 minutes to denature the RNA and any other possible contaminants. Reactions were stopped by adding 80 μL water, then 2x volumes (200 μL) of RNA Binding Buffer (Zymo), vortexing, and finally adding 3x volumes (300 μL) of 100% EtOH and vortexing. This binding reaction was purified over the Zymo column as described above and analyzed by gel electrophoresis as described below.

### RNA gel electrophoresis, blotting, and imaging

Blotting analysis of ManNAz-labeled RNA was performed conceptually similar to a Northern Blot with the following modifications. RNA purified, enriched, or enzymatically digested and conjugated to a DBCO-biotin reagent as a described above was lyophilized dry and subsequently resuspended in 15 μL df-GLBII with 1x SybrGold (Thermo Fisher Scientific). To denature, RNA was incubated at 55°C for 10 minutes and crashed on ice for 3 minutes. Samples were then loaded into a 1% agarose-formaldehyde denaturing gel (Northern Max Kit, Thermo Fisher Scientific) and electrophoresed at 110V for 45 minutes. Total RNA was then visualized in the gel using a UV gel imager. RNA transfer occurred as per the Northern Max protocol for 2 hours at 25°C, except 0.45 μm nitrocellulose membrane (NC, GE Life Sciences) was used. This is critical for downstream imaging as most positively charged nylon membranes have strong background in the infrared (IR) spectra. After transfer, RNA was crosslinked to the NC using UV-C light (0.18 J/cm^2^). NC membranes were then blocked with Odyssey Blocking Buffer, PBS (Li-Cor Biosciences) for 45 minutes at 25°C. Note that the blocking buffer made with TBS or PBS, both sold from Li-Cor Biosciences, work similarly for this step. After blocking, Streptavidin-IR800 (Li-Cor Biosciences) was diluted to 1:10,000 in Odyssey Blocking Buffer and stained the NC membrane for 30 minutes at 25°C. Excess streptavidin-IR800 was washed from the membranes by three, serial washes of 0.1% Tween-20 (Sigma) in 1x PBS for 5 minutes each at 25°C. NC membranes were briefly rinsed in 1x PBS to remove the Tween-20 before scanning on an Odyssey LiCor CLx scanner (Li-Cor Biosciences) with the software set to auto-detect the signal intensity for both the 700 and 800 channels. After scanning, images were quantified with the LiCor software (when appropriate) in the 800 channel and exported.

### DMB assay for sialic acid detection

Unless otherwise noted, all chemicals were supplied by Sigma. Native sialic acids on RNA or DNA were derivatized with 4,5-methylenedioxy-1,2-phenylenediamine dihydrochloride (DMB) and detected via reverse phase high-performance liquid chromatography (HPLC) according to established methods(*24*). In brief, RNA samples were lyophilized, and 100 μg (or otherwise noted in specific figures) of each sample was dissolved in 2 M acetic acid. Sialic acids were hydrolyzed by incubation at 80°C for 2 hours, and then cooled to room temperature before the addition of DMB buffer (7 mM DMB, 0.75 M β-mercaptoethanol, 18 mM Na_2_SO_4_, 1.4 M acetic acid). Derivatization was performed at 50°C for 2 hours. Following the addition of 0.2 M NaOH, samples were filtered through 10 kDa MWCO filters (Millipore) by centrifugation and stored in the dark at −20°C until use. Separation was performed via reverse phase HPLC using a Poroshell 120 EC-C18 column (Agilent) with a gradient of acetonitrile in water: T(0 minutes) 2%; T(2 minutes) 2%; T(5 minutes) 5%; T(25 minutes) 10%; T(30 minutes) 50%; T(31 minutes) 100%; T(40 minutes) 100%; T(41 minutes) 2%; T(45 minutes) 2%. DMB-derivatized sialic acids were detected by excitation at 373 nm and monitoring emission at 448 nm. Sialic acids standards included N-acetylneuraminic acid (Neu5Ac; Jülich Fine Chemicals), N-glycolylneuraminic acid (Neu5Gc; Carbosynth), 3-deoxy-D-glycero-D-galacto-2-nonulosonic acid (KDN; Carbosynth), and the Glyko Sialic Acid Reference Panel (Prozyme).

### Subcellular fractionation

#### Isolation of highly pure nuclei

Nuclei are intricately entwined with the ER, posing a challenge to biochemically separate nuclei cleanly from the ER without mixing. Gagnon et al.(*37*) describe a protocol which cleanly recovers mammalian nuclei after processing without significant residual ER membrane attached. We performed this protocol on adherent ManNAz-labeled HeLa cells without modification to the step-by-step instructions published. Due to the stringent isolation of the nuclei, some fraction of nuclei themselves lyse during the process, contaminating the non-nuclear fraction. Therefore, when examining the fractionation results of this protocol, we consider only the signal left in the nucleus. Signal in the supernatant will be partially mixed ER, Golgi, cytosol, some nuclei, as well as other cellular compartments. After fractionation as per the protocol, TRIzol was used to extract and process the RNA.

#### Isolation of cytosol and crude membrane fractions

The ProteoExtract® Native Membrane Protein Extraction Kit (EMD Millipore) was used as per the manufactures protocol on adherent ManNAz-labeled HeLa cells. This kit uses serial lysis steps: first to gently release soluble cytosol proteins and RNA and second to rupture membranous organelles such as the plasma membrane, Golgi, and ER. Because the lysis buffers are gentle, residual ER/Golgi are left on the nuclear fraction and thus analysis of samples generated from this kit was limited to the efficiently separated soluble cytosolic fractions compared to the membranous fractions. After fractionation as per the kit’s protocol, TRIzol was used to extract and process the RNA as described above.

### Membrane protection assay

Large scale crude membranes were isolated using the Plasma Membrane Protein Extraction Kit (ab65400, Abcam) following the manufactures protocol, stopping after producing the membrane pellet but before separating plasma membrane from others. Typically, 10x 15cm plates of 80% confluent 293T cells used for each biological replicate. Membranes were resuspended in 800 μL KPBS (136 mM KCl, 10 mM KH_2_PO_4_, pH 7.25 was adjusted with KOH (*42*)), 125 mM sucrose, and 2 mM MgCl_2_, split into 4 reactions, and incubated at 37°C for 1 hour with or without 0.1% Triton X-100 or 150 nM VC-Sia (homemade as per above). RNA was extracted with TRIzol and processed as described above for DMB analysis of sialic acid levels.

### Antibodies for western blotting

The following were used for blotting on nitrocellulose membranes at the indicated concentrations: 1:1000 GAPHD (A300-641A, Bethyl), 1:3000 β-tubulin (ab15568, Abcam), 1:5000 H3K4me3 (ab8580, Abcam), 1:1000 RPN1 (A305-026A, Bethyl), 1:1000 Sec63 (A305-084A, Bethyl). Appropriate secondary antibodies conjugated to LiCor IR dyes (Li-Cor Biosciences) and used at a final concentration of 0.1 ng/μL.

### Sucrose gradient fractionation of RNA

RNA used as input for sucrose gradient fractionation was previously extracted, PK treated, and clicked to DBCO-biotin as described above. RNA was sedimented through 15-30% sucrose gradients following McConkey’s method(*43*). Typically, 250-500 μg total RNA was lyophilized and then dissolved in 500 μL buffer containing 50 mM NaCl and 100 mM sodium acetate (pH 5.5). Linear 15-30% sucrose gradients were prepared in 1×3.5 inch polypropylene tubes (Beckman) using a BioComp 107 Gradient Master. Dissolved RNA was layered on top of pre-chilled gradients, which were then centrifuged using a SW32 Ti rotor at 80,000x g (25,000 rpm) for 18 hours in a Beckman Coulter Optima L70-K Ultracentrifuge at 4°C. Gradients were fractionated using a Brandel gradient fractionation system, collecting 0.75 mL fractions. Fractionated RNA was subsequently extracted from the sucrose solution using TRIzol as described above and analyzed by agarose gel electrophoresis or deep sequencing.

### Enrichment, deep sequencing, and analysis of ManNAz-labeled RNA

Two rounds of selection performed on RNA samples before sequence analysis to identify transcripts modified with ManNAz-containing glycans. Total RNA from ManNAz-labeled H9 or HeLa cells was extracted, purified, and conjugated to DBCO-biotin as described above. Biological duplicates, at the cell culture level (different passage number), were generated for the purposes of the sequencing experiments. The first enrichment was achieved by sucrose gradient fractionation; after centrifugation fractions containing small RNAs were pooled and TRIzol extracted. The second enrichment was achieved by selective affinity to streptavidin beads as previously published(*44*) with the following specific steps: 10 μL of MyOne C1 Streptavidin beads (Thermo Fisher Scientific), per reaction were blocked with 50 ng/μL glycogen (Thermo Fisher Scientific) in Biotin Wash Buffer (10 mM Tris HCl pH 7.5, 1 mM EDTA, 100 mM NaCl, 0.05% Tween-20) for 1 hour at 25°C. Biotinylated small RNAs from H9 and HeLa cells were thawed and 150 ng of each were saved for input library construction. Next, 25 μg of the biotinylated small RNAs were diluted in 750 μL Biotin Wash Buffer (final concentration of ~33 ng/μL) and mixed with the blocked MyOne C1 beads for 2 hours at 4°C. Beads were washed to remove non-bound RNAs: twice with 1 mL of ChIRP Wash Buffer (2x SSC, 0.5% SDS), twice with 1 mL of Biotin Wash Buffer, and twice with NT2 Buffer (50 mM Tris HCl, pH 7.5, 150 mM NaCl, 1 mM MgCl_2_, 0.005% NP-40), all at 25°C for 3 minutes each.

To construct deep sequencing libraries two approaches were taken using the same enzymes with different steps for the input(*45*) vs. bead-enriched(*46*) samples given that the latter were already conjugated to a bead-support.

#### Input libraries

The 150 ng of small RNAs isolated before MyOne C1 capture were lyophilized dry and then T4 PNK mix (2 μL 5x buffer (500 mM Tris HCl pH 6.8, 50 mM MgCl_2_, 50 mM DTT), 1 μL T4 PNK (NEB), 1 μL FastAP (Thermo Fischer Scientific), 0.5 μL SUPERaseIn, and 5.5 μL water) was added for 45 minutes at 37°C. Next, a pre-adenylated-3’linker was ligated by adding 3’Ligation Mix (1 μL of 3 μM L3-Bio_Linker(*5*), 1 μL RNA Ligase I (NEB), 1 μL 100 mM DTT, 1 μL 10x RNA Ligase Buffer (NEB) and 6 μL 50% PEG8000 (NEB)) to the T4 PNK reaction and incubating for 4 hours at 25°C. Unligated L3-Bio_Linker was digested by adding 2 μL of RecJ (NEB), 1.5 μL 5’ Deadenylase (NEB), 3 μL of 10x NEBuffer 1 (NEB) and incubating the reaction at 37°C for 60 minutes. Ligated RNA was purified with Zymo columns as described above and lyophilized dry. cDNA synthesis, enrichment of cDNA:RNA hybrids, cDNA elution, cDNA circularization, cDNA cleanup, first-step PCR, PAGE purification, and second-step PCR took place exactly as previously described(*6*).

#### Bead-enriched libraries

Washed MyOne C1 beads bounded to ManNAz-labeled small RNAs were processed as described before(*45*) with the following modifications. For the on-bead ligation step, a non-biotinylated 3’linker oligo was used (L3-Linker(*5*)) such that all RNAs captured on the beads would be included in the sequencing library. After completing the second-step PCR for both the input and bead-enriched samples, the dsDNA libraries were quantified on a High Sensitivity DNA Bioanalyzer chip (Agilent) and sequenced on a NextSeq 500 instrument (Illumina).

#### Data analysis

Sequencing data were processed largely as described previously with a pipeline designed to analyze infrared CLIP data(*6*). The specific version of the pipeline used in this work can be found here (https://github.com/ChangLab/FAST-iCLIP/tree/lite). Specifically, the raw reads were removed of PCR duplicates, adaptor sequences trimmed, mapped to a human genome reference (GCRh38) and custom sequence indexes of human repetitive RNAs (such as snRNAs and rRNAs), and finally reverse transcriptase (RT) stops were identified. We hypothesized that cDNA synthesis of unfragmented small RNAs used for the input samples would generate RT stops largely at the 5’ends of RNAs which would represent full length cDNA synthesis. In contrast, if the ManNAz-enrichment successfully isolated covalently modified RNAs, the RT may stop adjacent to the position where the RNA was modified, akin to the RT stopping adjacent to a UV-crosslinked nucleotide in irCLIP experiments(*6*). Enrichment of RNA families and specific RNA transcripts were performed by calculating relative proportions of unique RT stop counts in input and enriched libraries. Fractional nucleobase identity at RT stops in input and enriched libraries were used to calculate the enrichment of specific nucleobases in the snRNA and Y-RNA families.

### CRISPR/Cas9 knockout of Y5 and characterization

CRISPR gRNA sequences were designed using the CHOPCHOP online webtool (http://chopchop.cbu.uib.no/index.php) (*47*). Guides that flank the Y5 locus were selected (**table S4**). Corresponding oligos were ordered from IDT. Oligos were cloned into the Zhang lab generated Cas9 expressing pX458 guide RNA plasmid (Addgene) as previously described(*48*) using Gibson assembly reaction (NEB). Two sgRNAs flanking the human Y5 locus encoded in the pX458 plasmids were co-transfected using Lipofectamine 3000 (Thermo Fischer Scientific) according to manufactures guidelines in a 6-well format. Transfected cells were single cell sorted based on GFP expression into 96-well plates using BD influx cell sorter (Stanford FACS facility). Clonal cell lines were allowed to expand, and genomic DNA was isolated for sequenced based genotyping of targeted allele. For this, a 300–500 base-pair region that encompassed the gRNA-targeted site was amplified and the PCR product was Sanger sequenced. Clones with editing events causing large deletions were selected for subsequent experiments and KO loss of expression was confirmed by Northern blotting (below). To evaluate doubling time, 293 WT and KO cells were cultured as described above, initially seeding 20,000 cells per 12-well plate in triplicate. At 24-hour intervals cells were trypsinized and counted using a Countess II FL Automated Cell Counter (Thermo Fischer Scientific) instrument as per the manufacturer’s instructions.

### Small RNA Northern blotting

Detection of small RNAs was achieved by conventional Northern blotting and detection via radiolabeled locked-nucleic acids (LNAs). LNAs (Qiagen) complementary to the Y5 RNA or 5S rRNA (table S4) were ordered and 5’end labeled: 200 pmol LNA was added to 3 μL of T4 PNK (NEB), 7 μL 10x T4 PNK buffer, and 1 μL of ATP, [γ-^32^P]-3000 Ci/mmol 10 mCi/ml (γ-ATP, Perkin Elmer) in a 70 μL reaction. LNAs were incubated at 37°C for 3 hours after which free γ-ATP was purified away using Micro Bio-Spin 6 (Bio-Rad) columns as per the manufacture’s’ instructions. A 12% Urea-PAGE gel (National Diagnostics) was poured and pre-run at 10W for 15 minutes, after which 2 μg of total RNA from various cell types was separated by running the gel at 15W. After electrophoresis, RNA was transferred to HyBond N+ (GE Life Sciences) using a Semi-Dry transfer apparatus (Bio-Rad) with 0.5x Tris/Borate/EDTA (TBE, Thermo Fischer Scientific) buffer at a constant power of 18V for 90 minutes at 4°C. Next, RNA was crosslinked to the membrane, and pre-hybridized at 65°C for 60 minutes in 2 mL of PerfectHyb Plus (Sigma) buffer. Labeled LNA probes were then added to the PerfectHyb Plus buffer (typically 25% of the labeled LNA probe was used for any single membrane hybridization) and incubated at 65°C for 3-16 hours (no change in results with longer or shorter hybridizations). Membranes were rinsed 2x 2.5 mL of Low Stringency Northern Buffer (0.1% SDS, 2x SSC (Saline-sodium citrate)) and then washed at 37°C for 2x 5 minutes in 2.5 mL of High Stringency Northern Buffer (0.1% SDS, 0.5x SSC). Wash membranes were exposed to storage phosphor screens and finally imaged with a GE Typhoon 9410 scanner.

### Secondary Structure Folding of Y5 RNA

The human Y5 sequence was obtained (NR_001571.2) and secondary structure folded using the mFold web server(*49*) using default settings. A Vienna file (.b) was exported for the folded structure and visualized in VARNA(*50*).

